# Shared and modality-specific brain networks underlying predictive coding of temporal sequences

**DOI:** 10.1101/2025.07.16.665207

**Authors:** Leonardo Bonetti, Gemma Fernández-Rubio, Mattia Rosso, Francesco Carlomagno, Antonio Malvaso, Antonino Greco, Morten L. Kringelbach, Peter Vuust

## Abstract

Predictive coding posits that the brain continuously generates and updates internal models to anticipate incoming sensory input. While auditory and visual modalities have been studied independently in this context, direct comparisons using matched paradigms are scarce. Here, we employed magnetoencephalography (MEG) to investigate how the brain of 83 participants encodes and consciously recognises temporally unfolding sequences that acquire Gestalt-like structure over time, a feature rarely addressed in cross-modal research. Participants memorised matched auditory and visual sequences with coherent temporal structure and later identified whether test sequences were familiar or novel. Multivariate decoding revealed robust discrimination between the brain mechanisms underlying encoding and recognition of memorised and novel sequences, with sustained temporal generalisation in the auditory domain and time-specific responses in the visual domain. Using the BROAD-NESS pipeline, we identified modality-specific and supramodal brain networks. Auditory memory engaged auditory cortex, cingulate gyrus, and hippocampus, whereas visual memory involved orbitofrontal cortex and visual areas. Notably, both modalities recruited a shared network including hippocampus and medial cingulate during recognition. These findings provide compelling evidence for distinct and shared predictive learning mechanisms across sensory systems, advancing our understanding of how the brain integrates and evaluates temporally structured, Gestalt-like information.

## Introduction

The human brain organises information hierarchically, integrating both spatial and temporal dimensions to support perception and cognition. One influential theoretical framework explaining this process is predictive coding theory, which proposes that the brain continuously generates and updates predictions about incoming sensory input. By minimising the discrepancy between expected and actual stimuli, termed prediction error, the brain refines its internal models to optimise perception and behaviour (Feldman & Friston, 2010; Friston & Kiebel, 2009; Friston, 2010). This predictive framework is especially relevant for the visual and auditory systems, where the brain updates expectations in real time based on ongoing sensory input. In vision, predictive models help recognise objects, motion, and depth; in audition, they support the processing of speech and music by tracking temporal sequences and changes in frequency (Bendixen, 2014; Friston, 2010; Koelsch et al., 2019; Kveraga et al., 2007; Summerfield & Egner, 2009; Vuust et al., 2022).

Within the visual domain, neuroimaging research has extensively investigated how the brain processes static images (e.g., Carlson et al., 2013; Cichy et al., 2014; King et al., 2016). These studies provide support for the existence of hierarchical mechanisms that transfer information across the visual system, from the retina to the lateral geniculate gyrus in the thalamus, and then through the visual cortices to the inferior temporal cortex (DiCarlo et al., 2012; Jehee & Ballard, 2009; Spratling, 2010). However, visual perception is a dynamic process in real-world settings (Greco & Siegel, 2025), raising important questions about how the brain simultaneously represents multiple visual events presented over time. Building on this foundation, neurophysiological research has shifted towards the investigation of sequential information processing, particularly focusing on the conscious recognition of temporally structured sequences. These studies have typically used rapid serial visual presentation (RSVP) paradigms in which a target stimulus is briefly flashed within a sequence of non-target stimuli and recorded the brain activity related to processing each individual item with electroencephalography (EEG) or magnetoencephalography (MEG) (e.g., Grootswagers et al., 2019; King & Wyart, 2021; Marti & Dehaene, 2017; Potter et al., 2014). While such designs have yielded important insights into the brain dynamics of parallel and serial processing of visual stimuli, they did not factor in emergent perceptual principles described by Gestalt theory, which emphasises the brain’s tendency to integrate elements into coherent wholes rather than processing them as isolated components (Koffka, 2013; Wagemans et al., 2012). The auditory system offers instead a perceptual model that is ideally suited to the study of such emergent organisation due to the dynamic, time-evolving nature of auditory stimuli (Koelsch et al., 2019; Zatorre, 2003).

Auditory information is inherently sequential and conveys structured, holistic meaning, as seen in musical sequences (Koelsch et al., 2019). Accordingly, many auditory studies have used event-related potentials (ERPs) to investigate how the brain detects violations in predicted features of temporally structured sounds (Bonetti et al., 2017; Bonetti et al., 2018; Bonetti et al., 2021; Bonetti et al., 2022; Brattico et al., 2002; Conley et al., 1999; Greco et al., 2024b; Koelsch, 2009; Koelsch, 2011). Key ERP components include the N100, which reflects early auditory processing and short-term memory (Näätänen & Picton, 1987), the mismatch negativity (MMN), which signals unexpected deviations in acoustic input (Näätänen et al., 2007), and the error-related negativity (ERAN), which responds to syntactic violations in music and language, reflecting predictive mechanisms at higher levels (Koelsch, 2009). Building on these findings, we previously used MEG and musical memory paradigms to study how the brain encodes and recognises auditory sequences over time (Bonetti et al., 2023; Bonetti et al., 2024a; Bonetti et al., 2024b; Bonetti et al., 2024d; Bonetti et al., 2024e; Bonetti et al., 2025b; Costa et al., 2025; Fernández-Rubio et al., 2022a; Fernández-Rubio et al., 2022b; Fernández-Rubio et al., 2024; Herff et al., 2024; Nartallo-Kaluarachchi et al., 2025; Quiroga-Martinez et al., 2024; Serra et al., 2023). Under this approach, we identified a distributed network, from the auditory cortex to the medial cingulate, inferior temporal cortex, insula, frontal operculum, and hippocampus, that supports recognition of both familiar and novel musical sequences (Bonetti et al., 2023; Fernández-Rubio et al., 2022b; Fernández-Rubio et al., 2022c). More recently, we investigated the flow of information in the brain during such tasks using a refined experimental paradigm for musical recognition. We found that, for each tone in the sequence, brain activity originated in the auditory cortex and then propagated to the hippocampus, anterior cingulate, and medial cingulate gyrus, regions that are crucial both for confirming predictions and for detecting variations that elicit conscious prediction errors. Furthermore, we observed a backward flow of information from these regions to the auditory cortices (Bonetti et al., 2024d).

While our group has zoomed in on the predictive mechanisms of auditory perception and memory using musical sequences (e.g., Bonetti et al., 2024d; Bonetti et al., 2024e; Fernández-Rubio et al., 2022c), significant progress has been made in terms of the hierarchical dynamics supporting visual perception and memory for sequences of static images (e.g., King & Wyart, 2021; Marti & Dehaene, 2017). However, a combined, focalised neuroimaging approach that allows the direct comparison of visual and auditory sequences is still missing, raising several questions. How does the brain organize sequential visual inputs into a whole percept? How do the visual and auditory systems compare in this process, and where and when do their underlying mechanisms overlap or diverge? How does predictive coding interact with memory encoding and conscious recognition in both modalities? To address these questions, the present study examined the brain mechanisms underlying conscious encoding and recognition of temporally structured sequences in both the auditory and visual domains. Unlike previous studies, we created sequences of visual stimuli that maximize emergent features of Gestalt psychology, thus resembling musical sequences and allowing us to draw close parallels between the two modalities.

## Results

### Experimental design and behavioural results

Eighty-three participants completed an old/new recognition task while undergoing MEG recordings (Bonetti et al., 2024d). The experiment comprised two separate blocks targeting auditory and visual sensory modalities independently. Participants were initially presented 20 times with a sequence, either auditory or visual, and were instructed to memorise it thoroughly during the encoding phase. Auditory and visual sequences were matched in terms of their length, temporal structure, and contour (see Methods for details). During the subsequent recognition phase, participants were presented with 48 sequences: 24 were identical to the previously memorised sequence (M condition) and 24 were modified versions of the original (N condition). On each trial, participants indicated with a response pad whether the presented sequence was the one they had memorised or not.

**Figure 1.**
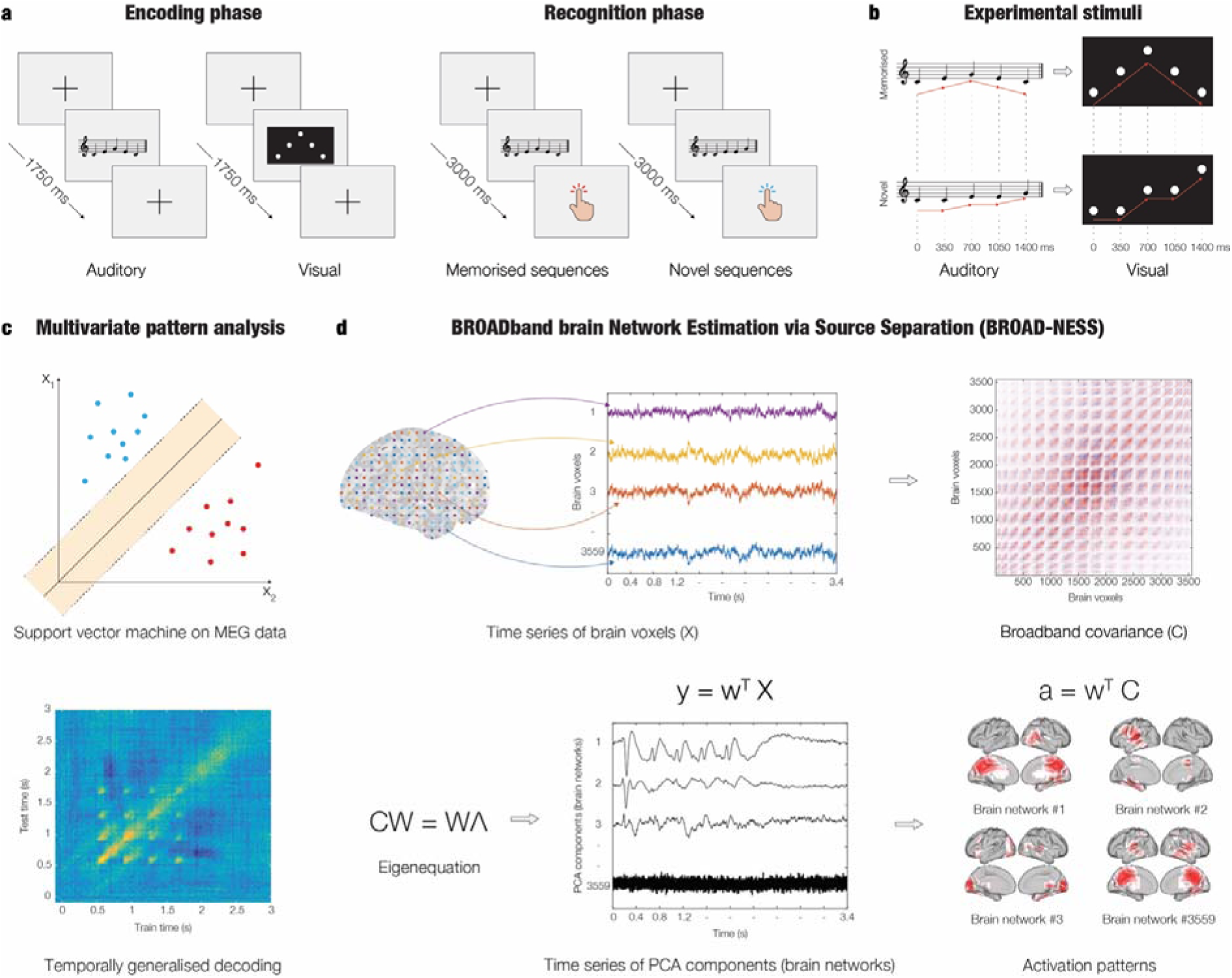
Experimental design, stimuli, and BROAD-NESS analytical pipeline. **a -** Participants (n = 83) underwent magnetoencephalography (MEG) recording while performing an old/new recognition task in separate auditory and visual blocks. Each block began with an encoding phase, during which a single sequence (auditory or visual) was repeated 20 times. This was followed by a recognition phase, where participants evaluated whether presented sequences were previously memorised (M) or novel (N) via button press. **b -** The auditory and visual sequences were structurally matched: each consisted of five items lasting 350 ms. The visual sequence mirrored the melodic contour of the auditory sequence. Novel sequences retained the first item but altered the remaining four, as detailed in the Methods. **c -** Multivariate pattern analysis (MVPA) was applied to the MEG data to determine whether neural activity could distinguish between experimental conditions and sensory modalities. A temporal generalisation approach was used, training a classifier at each time point and testing across all time points to identify stable patterns of neural activity. **d -** The BROAD-NESS (BROADband brain Network Estimation via Source Separation) analytical pipeline was used to extract large-scale brain network dynamics. MEG source-reconstructed signals from 3,559 brain voxels were used as input to Principal Component Analysis (PCA). Specifically, first the data was demeaned, then the covariance matrix was applied and finally the eigenvector problem was solved. This procedure identified principal components (PCs) representing large-scale brain networks, along with their associated time courses and brain voxel-level activation patterns.

Statistical analyses were conducted on behavioural data obtained from the MEG task. Specifically, four Wilcoxon signed-rank tests were performed to compare accuracy and reaction times between the M and N conditions, independently for each sensory modality. To control for multiple comparisons, a Bonferroni correction was applied, setting the adjusted significance threshold at ⍰ = .0125 (i.e., ⍰ = .05/4). The Wilcoxon signed-rank test showed that recognition accuracy was significantly higher for N than for M sequences in both the auditory (*z* = -2.70, *p* = .007, M = 22.94 ± 2.39, N = 23.35 ± 2.08) and visual tasks (*z* = -2.51, *p* = .012, M = 23.41 ± 2.07, N = 23.64 ± 1.44). Regarding reaction times, the Wilcoxon signed-rank test indicated that there was no significant difference between the two conditions in auditory recognition (*p* = .19). Conversely, reaction times were significantly lower in visual recognition of M compared to N sequences (*z* = -3.41, *p* < .001, M = 2150 ± 231 ms, N = 2226 ± 240 ms).

### Multivariate pattern analysis

We conducted several runs of multivariate pattern analysis (MVPA) to decode neural representations across distinct experimental conditions and sensory modalities (see Methods for further details). Seven pairwise comparisons were tested:

1. Auditory recognition previously memorised (ARM) versus auditory recognition novel (ARN)
2. Visual recognition previously memorised (VRM) versus visual recognition novel (VRN)
3. Auditory-visual recognition previously memorised (AVRM) versus auditory-visual recognition novel (AVRN)
4. Auditory encoding (AE) versus auditory recognition (AR)
5. Visual encoding (VE) versus visual recognition (VR)
6. Auditory encoding (AE) versus visual encoding (VE)
7. Auditory recognition previously memorised (ARM) versus visual recognition previously memorised (VRM)

As shown in **Figures 2**, **3** and **4**, each comparison yielded a temporal generalisation decoding matrix corrected with False Discovery Rate (FDR) for multiple comparisons that depicted the differentiation of neural responses between the two conditions. Overall, the results confirmed that the support vector machine (SVM) classifier successfully detected significant differences between M and N sequences across sensory modalities. Additionally, decoding reliably distinguished between encoding and recognition phases, and clearly separated auditory from visual processing, both during encoding and recognition.

**Figure 2.**
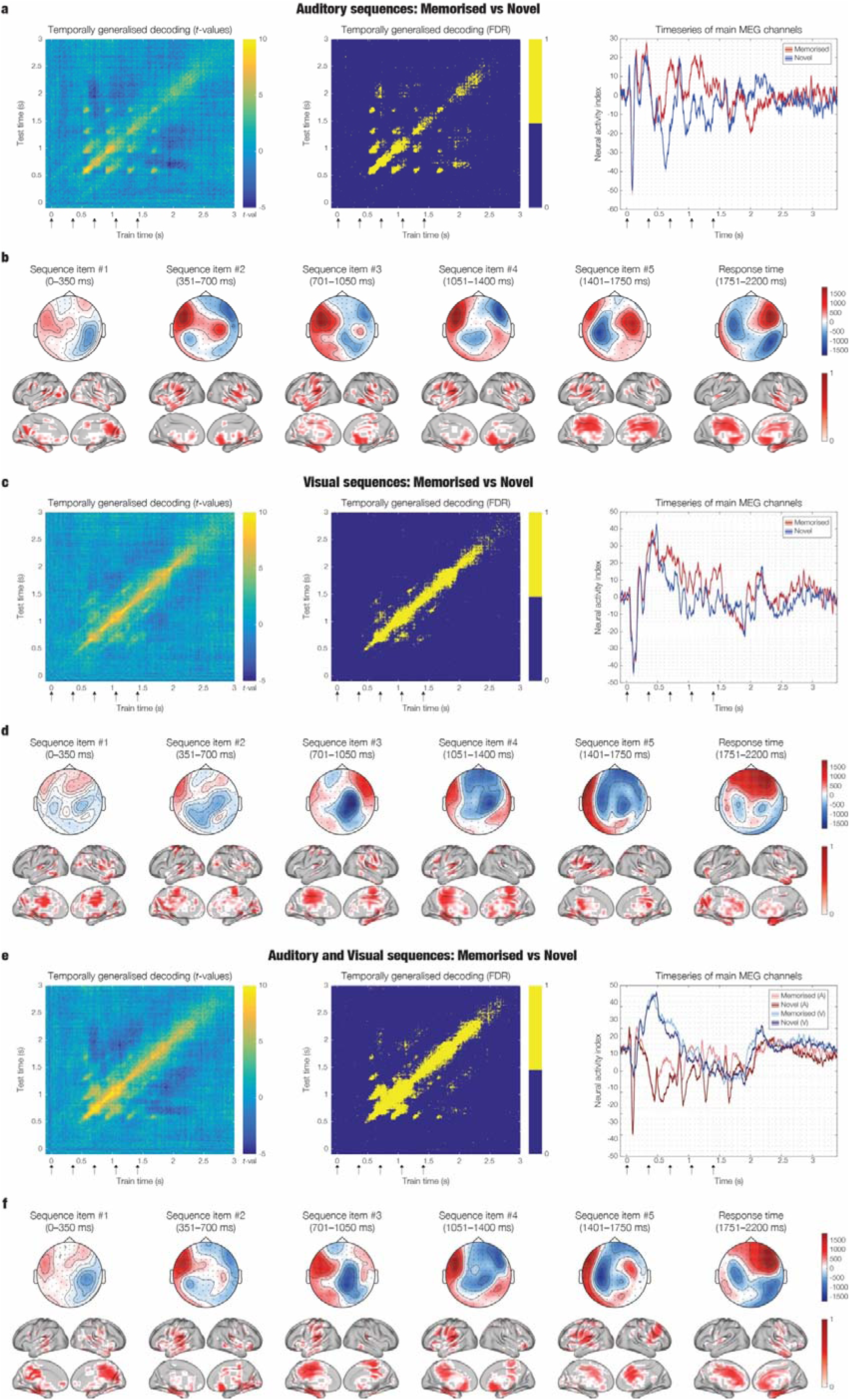
Decoding neural patterns underlying recognition of memorised versus novel sequences. **a -** Multivariate pattern analysis (MVPA) decoding of neural activity associated with the recognition of memorised (ARM) versus novel (ARN) musical sequences. The left panel shows t-values from a sign permutation test against chance level (50%) in a time-generalised decoding matrix. Arrows indicate the onset of each sound in the sequences. The middle panel highlights significant time-points where decoding accuracy exceeded chance after false discovery rate (FDR) correction (adjusted p < .002). The right panel presents the time series of decoding-relevant activity for M and N sequences, averaged over MEG channels contributing most strongly to the classifier (mean plus one standard deviation). **b -** Activation patterns derived from classifier weights, representing the relative contribution of each MEG sensor to decoding performance (unitless). The top row shows sensor-level topographies averaged over six relevant time windows corresponding to the five sequential sound onsets and the response period. These maps illustrate the spatial distribution of MEG sensor contributions to successful decoding at each key time point. The bottom row illustrates the source-level projections of the same activation patterns, reconstructed using beamforming, providing estimates of the cortical regions contributing most to the decoding of the recognition of memorised versus novel sequences. **c–f:** Equivalent decoding analyses for different stimulus modalities. **c, d:** Recognition of visual VRM versus VRN sequences (adjusted p < .001). **e, f:** Recognition of combined auditory and visual AVRM versus AVRN sequences (adjusted p < .012). All results represent participant-averaged data (n = 83); shaded areas in the time series plots indicate standard error of the mean.

**Figure 3.**
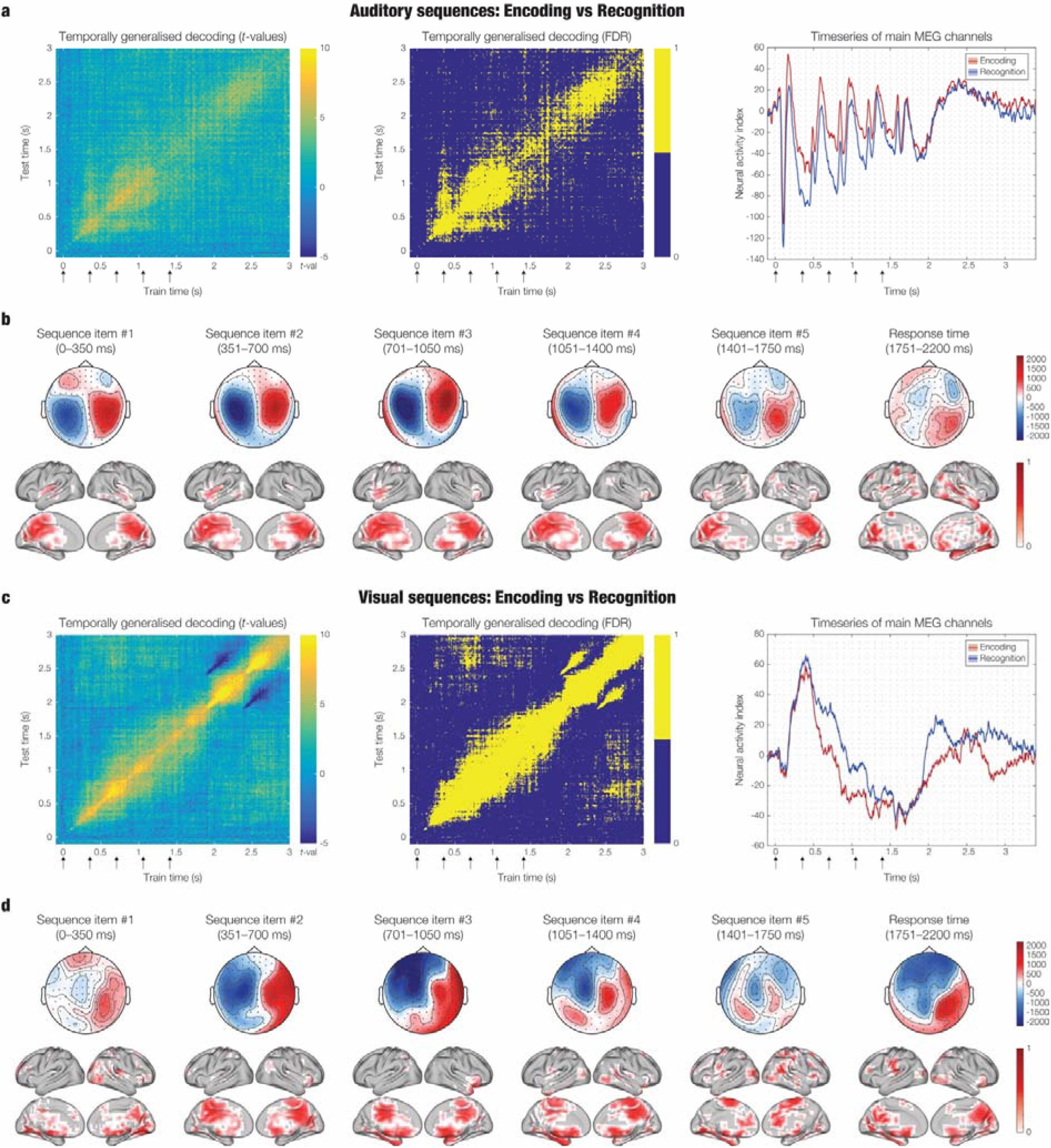
Decoding neural patterns underlying encoding versus recognition of sequences. **a -** Multivariate pattern analysis (MVPA) decoding of neural activity associated with the process of encoding (AE) versus recognising (AR) the same musical sequences. The left panel shows t-values from a sign permutation test against chance level (50%) in a time-generalised decoding matrix. Arrows indicate the onset of each sound in the sequences. The middle panel highlights significant time-points where decoding accuracy exceeded chance after false discovery rate (FDR) correction (adjusted p < .008). The right panel presents the time series of decoding-relevant activity for encoded and recognised sequences, averaged over MEG channels contributing most strongly to the classifier (mean plus one standard deviation). **b -** Activation patterns derived from classifier weights, representing the relative contribution of each MEG sensor to decoding performance (unitless). The top row shows sensor-level topographies averaged over six relevant time windows corresponding to the five sequential sound onsets and the response period. These maps illustrate the spatial distribution of MEG sensor contributions to successful decoding at each key time point. The bottom row illustrates the source-level projections of the same activation patterns, reconstructed using beamforming, providing estimates of the cortical regions contributing most to the decoding of encoded versus recognised sequences. **c**, **d:** Equivalent decoding analyses for encoding (VE) versus recognition (VR) of visual sequences (adjusted p < .013). All results represent participant-averaged data (n = 83); shaded areas in the time series plots indicate standard error of the mean.

**Figure 4.**
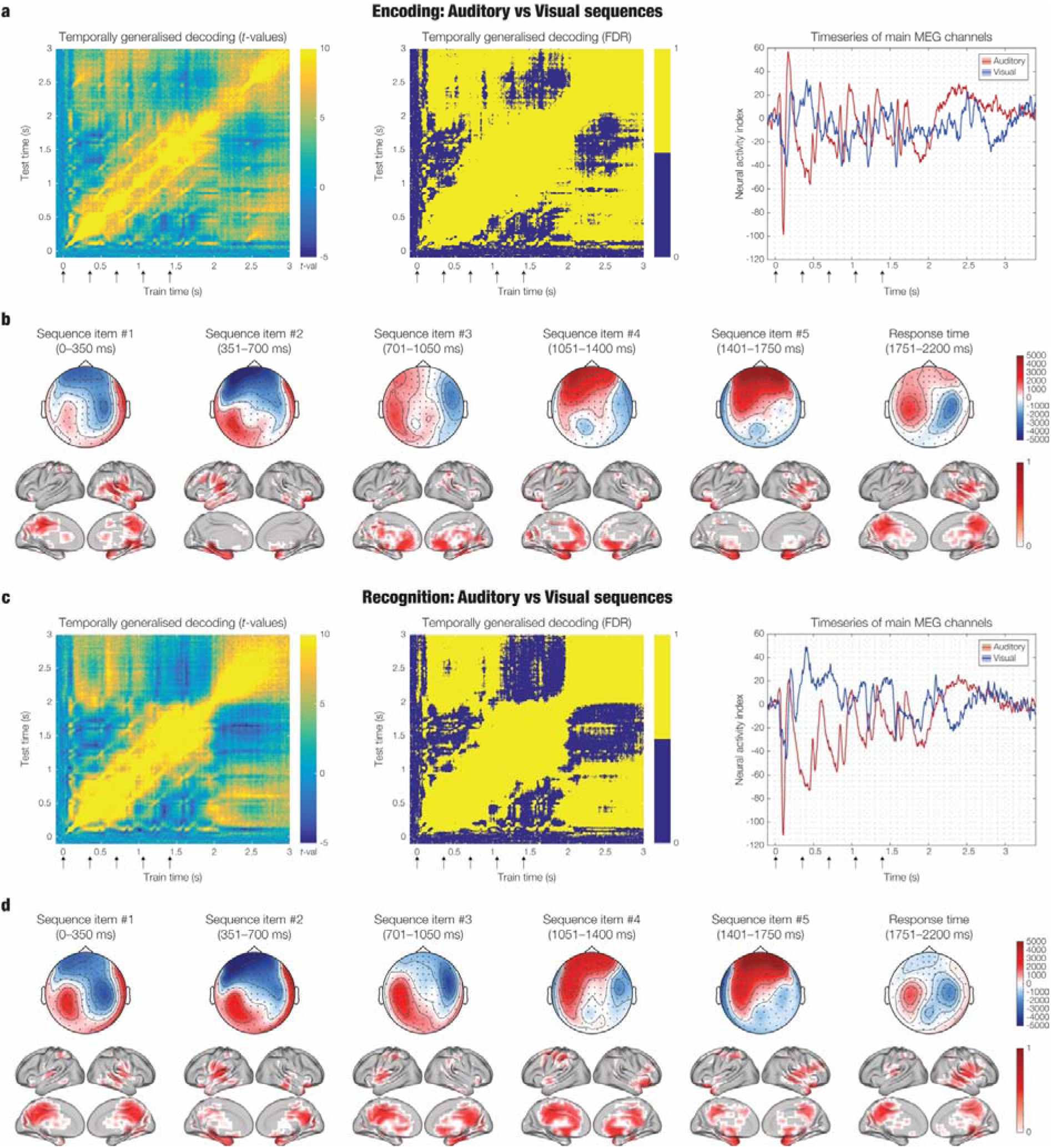
Decoding neural patterns underlying encoding and recognition of auditory versus visual sequences. **a -** Multivariate pattern analysis (MVPA) decoding of neural activity associated with the process of encoding auditory (AE) versus visual (VE) sequences. The left panel shows t-values from a sign permutation test against chance level (50%) in a time-generalised decoding matrix. Arrows indicate the onset of each item in the sequences. The middle panel highlights significant time-points where decoding accuracy exceeded chance after false discovery rate (FDR) correction (adjusted p < .041). The right panel presents the time series of decoding-relevant activity for encoded auditory and visual sequences, averaged over MEG channels contributing most strongly to the classifier (mean plus one standard deviation). **b -** Activation patterns derived from classifier weights, representing the relative contribution of each MEG sensor to decoding performance (unitless). The top row shows sensor-level topographies averaged over six relevant time windows corresponding to the five item onsets of the sequence and the response period. These maps illustrate the spatial distribution of MEG sensor contributions to successful decoding at each key time point. The bottom row illustrates the source-level projections of the same activation patterns, reconstructed using beamforming, providing estimates of the cortical regions contributing most to the decoding of encoded auditory versus visual sequences. **c**, **d:** Equivalent decoding analyses for recognition of auditory (AR) versus visual (VR) sequences (adjusted p < .039). All results represent participant-averaged data (n = 83); shaded areas in the time series plots indicate standard error of the mean.

More specifically (**Figure 2**), decoding ARM versus ARN revealed significant classification along the main diagonal, as well as temporally generalised effects, suggesting that each sound in the sequence elicited similar patterns of neural activity across time (*p* < .002, after false discovery rate [FDR] correction). Decoding VRM versus VRN produced a strong and temporally specific decoding pattern along the diagonal, with limited temporal generalisation, indicating that neural responses were more time-locked to individual visual items (*p* < .001, FDR-corrected). Decoding AVRM versus AVRN showed robust diagonal decoding and moderate temporal generalisation, intermediate results between the effects observed in ARM versus ARN and VRM versus VRN, reflecting the integration of auditory and visual information (p < .012, FDR-corrected).

Decoding AE versus AR (**Figure 3**) resulted in significant decoding along the diagonal and widespread temporal generalisation, pointing to a stable, sustained neural process distinguishing encoding from recognition (*p* < .008, FDR-corrected). Interestingly, this pattern differed from the generalisation seen in ARM versus ARN. While the latter showed temporally discrete generalisation for each sound, the AE versus AR comparison reflected a more continuous differentiation process over time. Decoding VE versus VR revealed strong decoding along the diagonal, but with limited temporal generalisation, suggesting that visual encoding and recognition may rely on temporally sharper, less sustained neural mechanisms (*p* < .013, FDR-corrected).

Finally, decoding AE versus VE (**Figure 4**) yielded widespread significant decoding effects, especially along the diagonal (*p* < .041, FDR-corrected), reflecting well-known differences in sensory processing between auditory and visual modalities. Similarly, decoding AR versus VR produced extensive decoding along the diagonal (*p* < .039, FDR-corrected). Notably, for both AE versus VE and AR versus VR, significant classification results extended beyond the stimulation window, indicating that modality-specific processes continued even after the stimulus offset.

To determine which MEG channels contributed most to the decoding algorithm, we computed activation patterns derived from the MVPA and averaged them for illustrative purposes across six time windows. These windows corresponded to the duration of each of the five items in the sequences, as well as the period from the end of the sequences to the participants’ average reaction time. To identify the brain sources associated with these MEG channels, we performed source reconstruction using a local-spheres forward model and a beamforming approach as the inverse solution, applied to an 8-mm grid comprising 3559 brain voxels. This analysis was conducted using magnetometers only. As detailed in the Methods section, we used the weights obtained from the beamforming algorithm to reconstruct the activation patterns derived from the MVPA. For visualisation purposes, these source-level patterns were also averaged across the same time windows used for the sensor-level data.

Figures 2, 3 and 4 show the contribution of each MEG channel and brain source to the decoding algorithm during the significant time windows.

### Brain networks underlying memory for temporal sequences

To investigate the brain networks involved in the encoding and recognition of auditory and visual sequences, we employed BROADband brain Network Estimation via Source Separation (BROAD-NESS), a framework that utilises Principal Component Analysis (PCA) to identify broadband brain networks from the reconstructed MEG time series (Bonetti et al., 2024c). PCA was applied to the 3559 brain voxel time series reconstructed from MEG recordings, to capture concurrent neural activity during the encoding and recognition of auditory and visual sequences, thereby revealing the brain networks underlying these processes.

This procedure, conducted independently for auditory and visual sequences, yielded three key outputs (Figure 5): (1) the variance explained by each principal component (PC), indicating the relative prominence of each network, (2) the spatial activation patterns of each PC, reflecting the network’s spatial extent, and (3) the activation time series for each brain network in every experimental condition.

**Figure 5.**
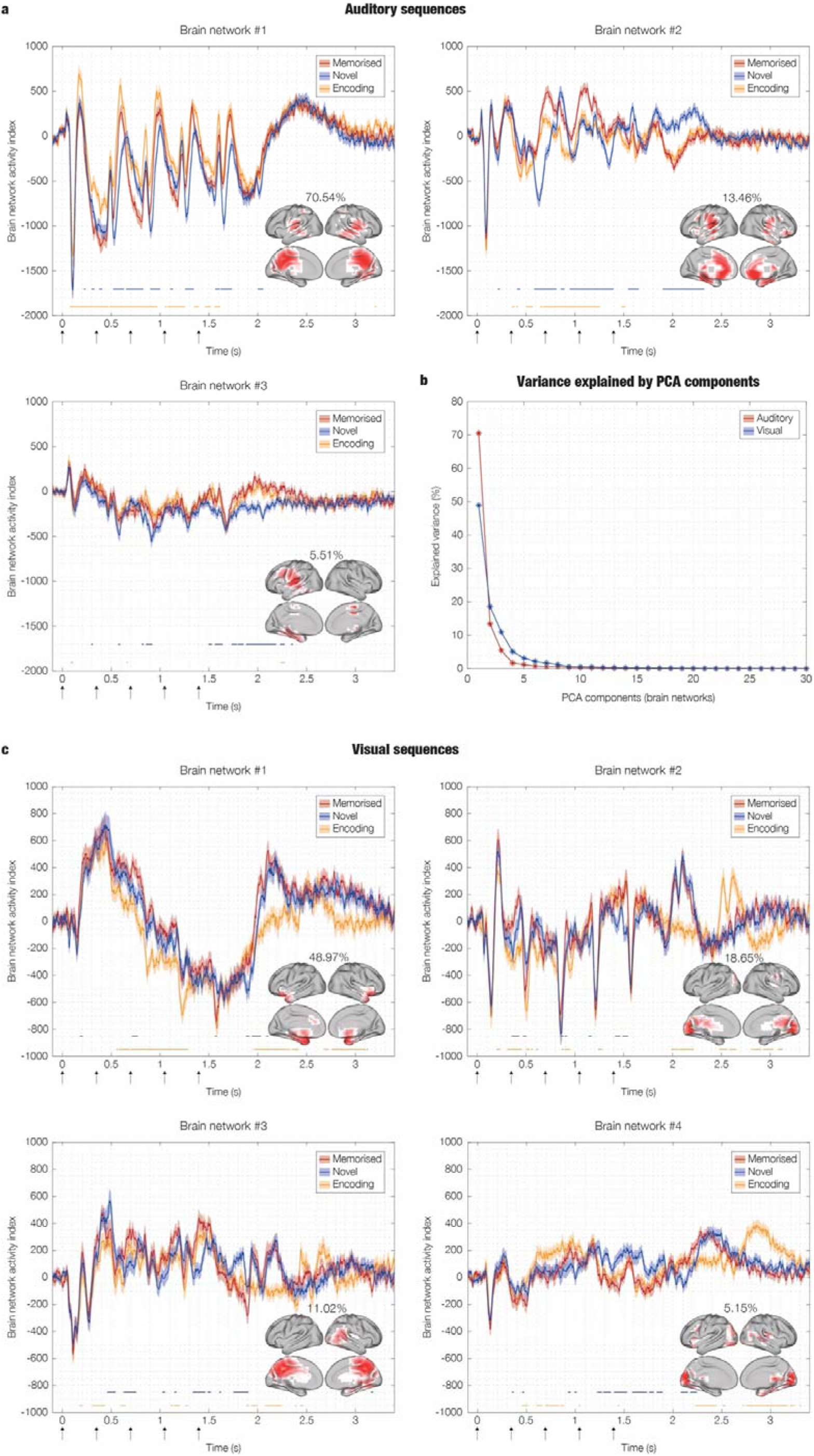
Brain networks underlying encoding and recognition of auditory and visual sequences. **a -** The figure displays the time series, variance explained and spatial activation patterns of the three brain networks explaining the highest variance in the auditory dataset. These networks were identified using principal component analysis (PCA) within the BROAD-NESS framework, computed on data averaged across conditions and participants. After PCA computation, the individual time series for each participant, brain network, and experimental condition were generated using PCA-derived weights, as described in the Methods section. These individual time series were then averaged across participants, as shown in the plots. Shaded areas represent standard errors. The brain templates illustrate the spatial extent of the networks, with dark red voxels contribute most strongly to the brain network and time series. Here, only voxels exceeding the mean by more than one standard deviation (in absolute terms) are shown. Orange and blue lines indicate the time windows during which significant differences were observed between the encoding and recognition of melodies, and between the recognition of M and N melodies, respectively. These contrasts were assessed using two-sided t-tests and corrected for multiple comparisons using cluster-based permutation tests. **b -** Variance explained by the principal components after PCA on the auditory and visual datasets. A similar proportion of variance was explained by three networks in the auditory dataset and by four networks in the visual dataset, suggesting a more modular organization involving a greater number of concurrently active memory networks in the visual system compared to the auditory system. **c -** Brain networks for the visual dataset, following the same depiction conventions as in **(a)**.

To determine the optimal number of PCs to retain, we applied the elbow rule in the context of the characteristic exponential decay of explained variance. This revealed three main brain networks for auditory sequences, explaining 70.54%, 13.46%, and 5.51% of the variance, respectively, and four brain networks for visual sequences, explaining 48.97%, 18.65%, 11.02%, and 5.15% of the variance, respectively (Figure 5b).

To compare brain network dynamics across experimental conditions, we statistically evaluated the network time series. Specifically, we performed two-sided t-tests at each time point for the following contrasts, separately for auditory and visual conditions:

1. Recognition of previously memorised sequences (M) vs encoding (E)
2. Recognition of previously memorised sequences (M) vs recognition of novel sequences (N)

To correct for multiple comparisons, we applied a cluster-based permutation test, following the method described by Maris and Oostenveld (2007), with a *p* value threshold of .001.

As shown in Figure 5a, the first auditory network comprised the auditory cortices and the medial cingulate gyrus. The second network included the auditory cortices, anterior cingulate gyrus, ventromedial prefrontal cortex, hippocampal regions, and the insula. The third network was primarily composed of the left auditory cortex and hippocampal regions. The time series of these networks revealed several significant time windows that survived cluster-based correction.

For the visual sequences, the first identified network encompassed bilateral orbitofrontal cortices extending into the anterior hippocampal regions. The second network was primarily localised to the occipital lobe, particularly the visual cortex. The third network included the medial cingulate gyrus and right hippocampal regions, while the fourth involved both the visual cortex and right hippocampus. Similar to the auditory results, the visual network time series exhibited several significant time windows that passed the cluster-based correction threshold. These are depicted in Figure 5c.

## Discussion

This study investigates how the brain supports predictive processing and memory for temporal sequences across auditory and visual modalities, aiming to uncover the shared and distinct neural mechanisms that underlie memory for sequences. Using matched auditory and visual paradigms, we examined how the brain encodes and recognises sequential information in both domains, extending previous work by focusing on consciously perceived, temporally extended sequences. Behaviourally, participants showed slightly higher accuracy for recognising novel compared to memorised sequences in both modalities, while reaction time differences emerged only in the visual domain. MVPA successfully decoded M versus N sequences in both auditory and visual tasks, as well as encoding versus recognition phases. Notably, decoding was also effective across modalities, revealing both shared and distinct neural representations. Finally, broadband source-level network analyses (i.e., BROAD-NESS) identified distinct spatiotemporal brain networks supporting encoding and recognition in each modality, with clear differences in the number, composition, and prominence of networks between auditory and visual systems.

Participants demonstrated slightly better accuracy for novel sequences than for memorised ones in both auditory and visual tasks, consistent with previous studies reporting superior memory for novel relative to familiar stimuli (i.e., novelty effect) (Kormi-Nouri et al., 2005; Tulving & Kroll, 1995). The absence of a significant difference in reaction times for the auditory task aligns with prior studies suggesting that recognition of auditory items, particularly musical ones, can occur relatively automatically (Koelsch, 2009). Conversely, the slower responses for visual sequences could reflect greater perceptual and attentional demands associated with temporally-unfolding visual information (VanRullen & Koch, 2003). While the novelty advantage appears robust in young and middle-aged adults, previous studies, including our own, have shown that this may diminish or even reverse with age. Older adults often display impaired recognition of novel sequences and greater reliance on familiarity (Belleville et al., 2011; Bonetti et al., 2024e; Bonetti et al., 2025a; Fernández-Rubio et al., 2024; Tanila et al., 1997), highlighting the cognitive cost of novelty processing and its susceptibility to age-related decline.

Our decoding results both replicate and extend previous research on predictive processing in the auditory domain. In line with earlier findings (Bonetti et al., 2024d; Bonetti et al., 2024e), we observed that M and N auditory sequences could be reliably distinguished from the point of divergence, beginning with the second sound. This supports predictive coding models, which posit that violations of expected sensory input elicit distinct neural responses (Heilbron & Chait, 2018). Auditory prediction violations have been well documented, both at an automatic level via MMN responses (Näätänen et al., 2007) and at a conscious level through recognition memory tasks such as the one employed here. The distributed neural pattern we observed, involving auditory cortices, insula, hippocampus, inferior temporal cortex, and critically the medial and anterior cingulate gyrus, closely mirrors previous findings in auditory memory and musical sequence processing (Bonetti et al., 2023; Bonetti et al., 2024d; Bonetti et al., 2024e; Koelsch et al., 2004; Koelsch, 2011). Crucially, our study advanced previous work by directly contrasting encoding and recognition phases of identical auditory sequences. Despite the perceptual similarity of the stimuli, MVPA revealed temporally sustained neural differences across the entire sequence, providing evidence that encoding and recognition engage overlapping yet functionally distinct brain mechanisms (Danker & Anderson, 2010; Rugg et al., 2012). This finding is consistent with predictive coding accounts, which suggest a transition from bottom-up sensory processing during encoding to top-down predictive reconstruction during recognition (Feldman & Friston, 2010; Friston & Kiebel, 2009; Friston, 2010). Among the regions driving this distinction, the medial cingulate gyrus, hippocampus, and insula emerged as key contributors, underlining their central role in memory-based auditory processing.

Our visual paradigm, involving the sequential appearance of dots to form recognisable shapes, enabled robust decoding of both encoding versus recognition phases and M versus N sequences during recognition. The strongest discriminative neural signals were consistently localised in the hippocampus and medial cingulate gyrus across both comparisons, suggesting that these regions play a key role in predictive memory for visual sequences. Notably, this extends previous work on visual memory, which has predominantly focused on static stimuli (Carlson et al., 2013; King et al., 2016), and relates to research on visual statistical learning, which is essential to map out higher-order structures (Greco et al., 2024a; Greco et al., 2025; Theeuwes et al., 2022; Turk-Browne et al., 2005). By using temporally structured, abstract visual sequences, our findings show that temporal integration processes are also integral to visual memory, supporting the existence of a dynamic, predictive component in this domain. This contributes novel evidence to the field of predictive coding in the visual system, showing that consciously recognised visual patterns also involve top-down, memory-based mechanisms.

Building on these findings, we then asked whether similar predictive memory processes might operate across sensory modalities. Indeed, cross-modal decoding revealed that neural responses to M versus N sequences generalised between auditory and visual tasks, confirming the presence of shared, supramodal mechanisms. These common neural signatures were most prominent in bilateral hippocampus, medial and anterior cingulate cortex, insula, and portions of the temporal lobe, regions frequently implicated in high-level memory and prediction. These results support the existence of content-general memory systems, likely rooted in medial temporal and prefrontal networks (Preston & Eichenbaum, 2013) and underscore the role of these areas in mediating predictive memory across distinct sensory inputs.

A notable feature of our results was the difference in temporal generalisation patterns. In the auditory domain, decoding between encoding and recognition, as well as M versus N comparisons, revealed temporally sustained differences, reflected in off-diagonal accuracy in the temporal generalisation matrices. This suggests that each item in the auditory sequence is processed in a stable manner throughout the musical sequence. In contrast, visual decoding was more temporally specific and lacked such generalisation (Dehaene et al., 2014). Each dot item evoked a distinct neural response, pointing to a more segmented or itemised processing strategy that is consistent with previous studies employing RSVP paradigms (e.g., King & Wyart, 2021; Marti & Dehaene, 2017). These modality differences may also explain why auditory sequences, such as melodies, are typically experienced and encoded in a more holistic or Gestalt-like manner, while visual sequences rely more on local compositional strategies (Snyder & Alain, 2007; Zacks & Swallow, 2007).

Our brain network analysis revealed both modality-specific and supramodal features of the neural architecture underlying memory-based prediction and recognition. In the auditory domain, we replicated and extended our previous work (Bonetti et al., 2023; Bonetti et al., 2024d; Bonetti et al., 2024e), identifying two principal networks. The first one comprised bilateral auditory cortices and medial cingulate gyrus. The second one included the auditory cortex, anterior cingulate, ventromedial prefrontal cortex, hippocampus, and insula. While the primary role of auditory cortices in sensory processing is well documented, the inclusion of cingulate cortex, hippocampus, and ventromedial prefrontal cortex is in line with models of predictive coding and auditory memory (Gaab et al., 2003; Wacongne et al., 2011). These regions appear to support the integration of top-down predictions with sensory input and the conscious evaluation of prediction outcomes. The medial cingulate-dominant network was more pronounced during recognition than encoding, implying a specific role in evaluating prediction accuracy. A third, weaker auditory network involving the left auditory cortex and hippocampus was also detected, potentially reflecting stimulus-specific mnemonic binding or residual signal interspersed with noise.

In contrast, visual sequence recognition involved a more modular and distributed network configuration, comprising four main functionally distinct networks. The most prominent network involved the bilateral orbitofrontal cortex (OFC) and anterior hippocampus, rather than visual sensory cortices. This differs markedly from the auditory domain, where early sensory areas dominated. The OFC has been implicated in abstract outcome evaluation, predictive inference, and memory-based decision-making (Kringelbach, 2005; Rolls, 2004; Schoenbaum et al., 2009). Its engagement likely reflects the need for higher-order integration and evaluation of novel visual configurations. The second visual network, centred on occipital cortices, likely supports basic visual encoding and perceptual analysis, consistent with models of hierarchical visual processing (Grill-Spector & Malach, 2004; Ungerleider & Haxby, 1994). However, its limited condition sensitivity implies a largely bottom-up function. The third network, including medial and posterior cingulate gyrus, right hippocampus, lingual and fusiform gyri, and posterior inferior temporal regions, has been frequently linked to visual memory retrieval (Kozlovskiy et al., 2012; Rolls, 2019). A fourth, minor network may represent secondary or compensatory processes, or a combination of signal and residual noise. Its presence nonetheless suggests overlapping subnetworks may assist recognition of unfamiliar, abstract visual stimuli.

Together, these findings suggest modality-specific organisational principles: auditory predictive memory involves integrated sensory-cognitive networks, whereas visual recognition employs a more compartmentalised, modular architecture. This may reflect differences in temporal continuity and the inherent structure of auditory versus visual inputs. In evolutionary terms, the two⍰dimensional, spatially correlated structure of light has driven the occipital cortex to develop retinotopic architectures with center–surround and orientation⍰selective receptive fields that decorrelate the 1/f statistics of natural images, maximizing spatial information transmission (Barlow, 1961; Bolanos et al., 2024; Froudarakis et al., 2014). By contrast, the one⍰dimensional, temporally unfolding nature of sound has shaped the temporal cortex into a tonotopic gradient endowed with spectrotemporal receptive fields finely tuned to the joint spectral and temporal modulations of natural acoustic environments, thereby optimizing high⍰fidelity temporal tracking (Francl & McDermott, 2022; Lewicki, 2002). However, parallels were evident, particularly between the second auditory and third visual networks, since both featured prominently the hippocampus, medial cingulate cortex, and inferior temporal regions. This indicates a shared supramodal core supporting recognition and predictive evaluation.

A key strength of this study lies in the matched temporal structure of visual and auditory stimuli, allowing a rigorous cross-modal comparison of sequence recognition. However, this approach is not without limitations. Although we carefully matched the temporal progression of auditory tones and visual dots, complete equivalence between modalities is unattainable due to inherent perceptual differences. Future research should examine how these disparities affect predictive processing, perhaps using abstract or synthetic stimuli that blur sensory boundaries. Further studies might also explore audiovisual integration involving naturalistic stimuli to assess how predictive coding operates when cues from multiple modalities must be integrated for recognition. Despite these limitations, our study offers a valuable step towards understanding the neural basis of prediction and memory across sensory systems.

In conclusion, this study provides novel insights into the brain mechanisms that support predictive memory and conscious recognition of temporally structured sequences in the auditory and visual domains. Through MEG decoding and network analysis, we identified both shared and modality-specific neural signatures. Our findings support a predictive coding account of sequence recognition across modalities while also revealing key differences in network organisation between sensory systems. These results contribute to a more unified understanding of how the brain anticipates, encodes, and consciously recognises complex temporal patterns.

## Methods

### Participants

The participant sample comprised 83 healthy volunteers (33 males and 50 females [sex as a biological attribute, self-reported]) aged between 19 and 63 years (mean age: 28.76 ± 8.06 years). The sample was recruited in Denmark, with participants originating from Western countries. Information regarding participants’ gender was not collected, as this lay outside the scope of the study. Their educational backgrounds were largely homogeneous: 73 participants either held a university degree (*n* = 54) or were enrolled as university students (*n* = 19). Of the remaining ten, 5 had obtained a professional qualification following secondary education, while the other 5 held a secondary school diploma.

The study received ethical approval from the Institutional Review Board (IRB) of Aarhus University (case number: DNC-IRB-2020-006). All experimental procedures adhered to the Declaration of Helsinki – Ethical Principles for Medical Research. Informed consent was obtained from all participants prior to the commencement of the experimental procedures, and all received compensation for their participation.

### Experimental stimuli and design

In this study, we employed an old/new auditory recognition paradigm during magnetoencephalography (MEG) recordings (Bonetti et al., 2023; Bonetti et al., 2024d; Bonetti et al., 2024e; Fernández-Rubio et al., 2022b; Fernández-Rubio et al., 2022c). The task was implemented across two distinct experimental blocks, involving different sensory modalities: auditory and visual. In each case, participants were initially presented with either an auditory or a visual sequence that was repeated 20 times and were instructed to memorise it as thoroughly as possible (encoding phase). Subsequently, they were presented with either the same sequence (previously memorised [M], 24 trials) or a modified version of the original sequence (novel [N], 24 trials) and were required to indicate, for each trial, whether the sequence matched the one they had memorised or not using a response pad (recognition phase). For the auditory condition, the sequence comprised five sounds, each lasting 350 ms, as illustrated in Figure 1. The visual sequence consisted of white dots that appeared sequentially on a black background. Each new dot was presented every 350 ms, forming a simple geometrical shape over time. The temporal structure of the visual sequences mirrored that of the auditory ones, and the overall contour, ascending or descending, was designed to match the pitch progression in the auditory sequences (Figure 1). In both modalities, variations were constructed by keeping the first item (sound or dot) constant and altering the subsequent four items. MIDI versions of the auditory sequences were created using Finale (version 26, MakeMusic, Boulder, CO). Both auditory and visual stimuli were presented using PsychoPy v3.0.

### Data acquisition

MEG recordings were conducted in a magnetically shielded room at Aarhus University Hospital (AUH), Aarhus, Denmark, using an Elekta Neuromag TRIUX system equipped with 306 channels (Elekta Neuromag, Helsinki, Finland). Data were sampled at 1000 Hz and analogue-filtered within the 0.1–330 Hz range. Prior to recording, each participant’s head shape and the positions of four Head Position Indicator (HPI) coils were digitised relative to three anatomical landmarks using a 3D digitiser (Polhemus Fastrak, Colchester, VT, USA). These data were later used to co-register the MEG recordings with individual anatomical magnetic resonance imaging (MRI) scans. Throughout the MEG session, the HPI coils continuously tracked the head position, enabling post hoc correction for head movement. To monitor physiological artefacts, two sets of bipolar electrodes were used to record electrocardiographic (ECG) and electrooculographic (EOG) activity, allowing for subsequent artefact removal during data preprocessing.

Structural MRI scans were acquired on a CE-approved 3T Siemens scanner at AUH. T1-weighted images (MPRAGE with fat saturation) were collected with a spatial resolution of 1.0 × 1.0 × 1.0 mm³. Sequence parameters were as follows: echo time (TE) = 2.61 ms; repetition time (TR) = 2300 ms; reconstructed matrix size = 256 × 256; echo spacing = 7.6 ms; bandwidth = 290 Hz/Px.

MEG and MRI data were acquired on two separate days.

### Behavioural data

We obtained behavioural data consisting of the number of correctly identified trials and the corresponding reaction times from the experimental task conducted during the MEG recordings, separately for the auditory and visual sequences. As the data did not follow a normal distribution, we employed four independent Wilcoxon signed-rank tests to evaluate whether accuracy and reaction times differed between the M and N conditions, independently for each sensory modality (auditory and visual). To account for multiple comparisons, we applied a Bonferroni correction, resulting in a significance threshold of ⍰ = .0125 (i.e., ⍰ = .05/4).

### MEG data pre-processing

Raw MEG sensor data (204 planar gradiometers and 102 magnetometers) were initially pre-processed using MaxFilter 70 (version 2.2.15) to attenuate external noise and environmental interferences. Signal Space Separation (SSS) was applied with the following parameters: spatiotemporal SSS, downsampling from 1000 Hz to 250 Hz, and movement compensation using continuous Head Position Indicator (cHPI) coils (default step size: 10 ms). The correlation threshold between inner and outer subspaces for rejecting overlapping signals during spatiotemporal SSS was set to 0.98.

We downsampled the data by a factor of four, given that MEG is relatively insensitive to high-gamma frequencies (e.g., >100 Hz) and exhibits higher signal fidelity in lower frequency bands (0.1–60 Hz), which were the focus of this study (Gross et al., 2013; Hansen et al., 2010). Furthermore, our interest lay in event-related broadband brain responses associated with memory recognition, which are typically expressed in lower frequency ranges (Albouy et al., 2017; Bonetti et al., 2023). This commonly adopted downsampling approach substantially reduced data volume, facilitating computational efficiency without compromising data quality.

The data were subsequently converted into Statistical Parametric Mapping (SPM) format and further processed in MATLAB (MathWorks, Natick, MA, USA) using a combination of in-house scripts (LBPD, https://github.com/leonardob92/LBPD-1.0.git) and the Oxford Centre for Human Brain Activity (OHBA) Software Library (OSL) (https://ohba-analysis.github.io/osl-docs/) (Woolrich et al., 2011). OSL builds upon established toolboxes including FieldTrip (Oostenveld et al., 2011), FSL (Woolrich et al., 2009), and SPM (Penny et al., 2011). Continuous MEG recordings were visually inspected using the OSLview tool to identify and remove large artefacts. The excluded data comprised less than 0.1% of the total dataset. Subsequently, independent component analysis (ICA), as implemented in OSL, was used to identify and remove components associated with eye blinks and cardiac artefacts (Mantini et al., 2011). The procedure involved decomposing the MEG signal into statistically independent components, correlating these with electrooculogram (EOG) and electrocardiogram (ECG) signals, and identifying components whose correlation coefficients exceeded three times those of the remaining components. These were further validated through visual inspection of their topographical distributions to confirm typical artefactual patterns. Components meeting both criteria were rejected, and the MEG signal was reconstructed from the remaining components. Finally, MEG data were epoched into 48 trials (24 M and 24 N), independently for auditory and visual experimental blocks. Each trial included a 100 ms pre-stimulus baseline period. Baseline correction was performed by subtracting the mean pre-stimulus signal from the post-stimulus data.

### Multivariate pattern analysis (decoding) – MEG sensors

We conducted several runs of multivariate pattern analysis (MVPA) to decode neural representations across different experimental conditions and blocks. Specifically, we tested the following seven pairwise comparisons:

1. Auditory recognition previously memorised (ARM) versus visual recognition novel (ARN)
2. Visual recognition previously memorised (VRM) versus visual recognition novel (VRN)
3. Auditory-visual recognition previously memorised (AVRM) versus auditory-visual recognition novel (AVRN)
4. Auditory encoding (AE) versus auditory recognition (AR)
5. Visual encoding (VE) versus visual recognition (VR)
6. Auditory encoding (AE) versus visual encoding (VE)
7. Auditory recognition previously memorised (ARM) versus visual recognition previously memorised (VRM)

This set of analyses was chosen as an initial step, since decoding leverages the entire dataset and does not rely on predefined assumptions regarding the selection of MEG channels or time points. Support Vector Machines (SVMs) (Steinwart & Christmann, 2008) were employed as classifiers, with independent analyses conducted for each participant and decoding analysis. MEG data were arranged into a three-dimensional matrix (channels × time points × trials) and entered into the SVM algorithm. To mitigate overfitting, we implemented a leave-one-out cross-validation procedure. Specifically, the dataset was partitioned into N subsets (with N = 4), where N – 1 subsets were used for training and the remaining subset for testing. This process was repeated so that each subset served as the testing set once. For each time point, decoding accuracy was computed. This procedure was repeated 100 times, each time randomly reassigning trials to training and testing subsets. The resulting decoding accuracy time series were then averaged across permutations to produce a final classifier performance curve for each participant. In addition to decoding accuracy, the classifier also produced weights and activation patterns, which indicate the MEG channels contributing most significantly to the classification (Haufe et al., 2014). To examine the temporal generalisability of neural patterns differentiating each condition pair, we also performed temporal generalisation analysis. In this analysis, the classifier trained on data from a single time point was tested on all time points, producing a two-dimensional matrix that reflects how stable and recurrent neural patterns are across time (Cichy et al., 2014). Statistical significance of decoding accuracy was evaluated using a signed permutation test against chance level (50%) for each time point. To control for multiple comparisons, we applied false discovery rate (FDR) correction (α = .05).

### Source reconstruction

Magnetoencephalography (MEG) is a powerful technique for capturing whole-brain neural dynamics with exceptional temporal resolution. However, to interpret the neural sources underlying these signals, spatial information is crucial. Since MEG records neural activity from outside the head, identifying the precise brain regions responsible for the recorded signals involves solving the so-called inverse problem. To address this, we employed beamforming algorithms (Brookes et al., 2007; Hillebrand & Barnes, 2005; Huang et al., 1999), implemented using a combination of in-house-developed code and open-source tools from the OSL, SPM, and FieldTrip toolboxes. The source reconstruction pipeline consisted of two main steps: (1) construction of the forward model and (2) computation of the inverse solution via beamforming.

A single-shell forward model was constructed for each participant using an 8-mm grid. This model assumes that each brain source can be represented as an active dipole and predicts how the signal from such a dipole would be detected by MEG sensors. MEG data were co-registered with individual T1-weighted MRI scans using anatomical landmarks captured via a 3D digitizer. In cases where individual MRI data were not available, a standard MNI152-T1 template (8-mm resolution) was used to construct the head model. Beamforming was then used as the inverse model to estimate the spatial sources of the recorded brain signals. Beamforming involves calculating spatial filters (i.e., sets of weights) that isolate the contribution of each potential dipole (brain source) to the MEG signal. This process was performed at each time point in the data, allowing for time-resolved source reconstruction. The signal recorded by MEG sensors at a given time *t*, denoted as *B(t)*, can be described by the following Eq.:

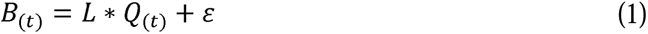

Here, *L* is the leadfield matrix (the forward model), *Q(t)* represents the dipole activity matrix at time *t*, and ε is a noise term (Huang et al., 1999). The inverse problem consists of estimating *Q(t)* from *B(t)*. In beamforming, this is achieved by computing a set of spatial weights *W*, such that the estimated dipole activity *q(t)* is given by:

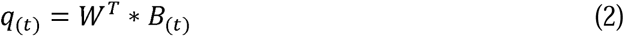

Where *W^T^* denotes the transpose of the weight matrix. To compute *W*, we used the covariance matrix *C* of the MEG channels, along with the leadfield matrix. Importantly, *C* was calculated by concatenating all relevant trials and experimental blocks, depending on the specific decoding contrast for which source reconstruction was performed. For instance, if a decoding analysis compared auditory and visual conditions, then both sets of trials were used to compute *C*. Conversely, when only auditory or visual data were involved, *C* was computed separately for each modality. This step was critical to avoid reconstruction bias. In fact, if the spatial filters are computed based only on one modality, they might not accurately reflect differences arising in cross-modal comparisons. In relation to the network analyses described in the subsequent section of the Methods, *C* was calculated independently for each sensory modality, as the network analyses were conducted accordingly.

Back to the Eq. (2), for each dipole *q*, the weights *W(q)* were computed using the following Eq. (3):

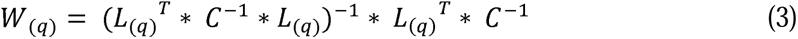

The leadfield model *L* was computed for the three principal orientations of each dipole (Nolte, 2003). Prior to calculating the weights, to simplify the beamforming output the dipole orientations were reduced to a single one using the singular value decomposition (SVD) algorithm applied to the matrix product described in Eq. (4).

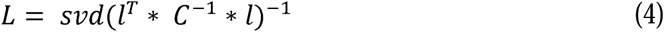

Here, *l* refers to the leadfield matrix containing all three orientations, and *L* represents the resolved single-orientation leadfield used in Eq. (3). Once the weights were computed, they were normalised to compensate for potential biases towards sources closer to the centre of the head (Luckhoo et al., 2014). These normalised weights were then applied in different ways, depending on the purpose of the source reconstruction.

First, we used the weights to project the activation patterns derived from the decoding analyses into source space. Specifically, the decoding-based sensor-space activation patterns (equivalent to matrix *B* in Eq. 2) were multiplied by the beamformer weights to generate 3559 source-resolved activation maps over time. For illustrative purposes, these reconstructed activation maps were averaged across six successive time windows, each corresponding to an item in the presented sequences and to the period during which participants made their recognition decisions. The first five windows were each 350 ms in duration, while the sixth was longer, defined as follows: (1) 0.001–0.350 s, (2) 0.351–0.700 s, (3) 0.701–1.050 s, (4) 1.051–1.400 s, (5) 1.401–1.750 s, and (6) 1.751–2.200 s.

Second, the normalised weights were applied to the averaged neural activity across trials, independently for each time point and experimental condition, yielding a time series for each of the 3559 brain sources (Brookes et al., 2007; Luckhoo et al., 2014), as described in Eq. (2). We used these source-level time series for subsequent functional network analyses, which are described in the following section.

### BROADband brain Network Estimation via Source Separation (BROAD-NESS)

To investigate the brain networks underlying encoding and recognition of auditory and visual sequences, we employed BROAD-NESS (Bonetti et al., 2024c), a framework that utilises Principal Component Analysis (PCA) to identify broadband brain networks from the reconstructed MEG time series. PCA is a dimensionality reduction technique that transforms high-dimensional data into a set of new variables, called principal components (PCs), which account for the largest variance in the data. This process is achieved by computing the eigenvectors and eigenvalues of the data’s covariance matrix and projecting the data onto the directions of maximum variance. PCA simplifies complex datasets while retaining the most important patterns. In this study, PCA was applied to the dense set of MEG-reconstructed brain data at the voxel level, specifically to the 3559 brain voxel time series, averaged over participants after adjusting the polarity of the time series reconstructed for each brain voxel, as previously done in analogous studies (Bonetti et al., 2024c; Bonetti et al., 2024d). The objective was not only to reduce dimensionality but also to identify brain networks (i.e., the PCs) that best captured the concurrent neural activity during encoding and recognition of auditory and visual sequences.

In short, PCA was computed as follows. The data was centred by subtracting the mean of each brain voxel timeseries from the dataset *X*, as shown by Eq. (5):

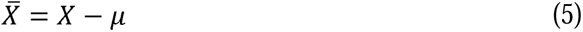

where µ is the mean vector of *X*.

The covariance matrix *C* was computed on the centred data, as shown in Eq. (6):

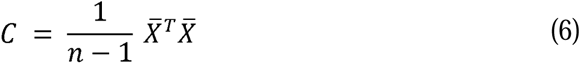

where *n* is the number of data points.

The eigenvalue Eq. for the covariance matrix was solved to find eigenvectors and eigenvalues, Eq. (7):

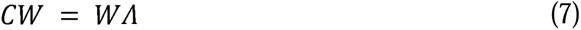

where *W* is the matrix of eigenvectors and Λ is the diagonal matrix of eigenvalues, which indicate the amount of variance explained by each component. Here, the components (brain networks) that explained significantly more variance than the others were selected, following the elbow rule for determining the number of PCs in the context of an exponential decay in explained variance. This process was done independently for the auditory and visual blocks.

Afterwards, for each brain network, the associated eigenvector *w* was used to compute its activation time series *y*. This was done by multiplying the transposed eigenvector by the original data *X*, as shown in Eq. (8):

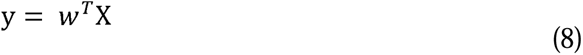

In addition, we computed the spatial projection of the components (spatial activation patterns *a*) in brain voxel space. We calculated this by multiplying the weights of the analysis (the eigenvectors w) by the covariance matrix *C*, as described by Eq. (9):

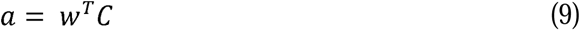

To be noted, due to the orthogonality of the PCs, the relative contribution of each brain voxel to the network is identical whether using the direct eigenvectors *w* or the computed spatial activation patterns *a*.

After computing the PCs and the corresponding network time series for each participant and experimental condition, we performed statistical tests to compare brain activity during different experimental conditions. Specifically, we performed a two-sided t-test for each time point, comparing the following contrasts independently for both auditory and visual experimental blocks:

1. Recognition previously memorised (M) versus encoding (E)
2. Recognition previously memorised (M) versus novel (N) sequences

Finally, to correct for multiple comparisons, we applied a cluster-based permutation test, as outlined by Maris and Oostenveld (2007). The procedure involved several steps. Initially, we computed statistical tests on the original data as described above. We then identified clusters of significant time points by applying a threshold of α = .05 to the statistical results. Next, we performed 1000 permutations, in which the labels of the two experimental conditions were shuffled for each participant, and the statistics were recalculated for each permutation, yielding a reference distribution of the significant cluster sizes (threshold of α = .05) that were obtained from all permutations. Clusters in the original data were considered significant if their sizes exceeded the maximum cluster size observed in the permuted data at least 99.9% of the time, which corresponds to a *p* value threshold of .001.

## Data availability

The data related to the experiment is available upon request.

## Code availability

The code is available at the following links: https://github.com/leonardob92/LBPD-1.0.git and https://github.com/leonardob92/MemoryPredictiveCoding_AuditoryVisualSequences.git.

## Acknowledgements

The Center for Music in the Brain (MIB) is funded by the Danish National Research Foundation (project number DNRF117). Additionally, we thank the Fundación Mutua Madrileña for the economic support provided to author GFR.

LB is supported by Sapere Aude: Independent Research Fund Denmark (DFF) Research Leader (grant ID: 10.46540/5253-00003B), Lundbeck Foundation (TalentPrize2022), Carlsberg Foundation (CF20-0239), Center for Music in the Brain, Linacre College of the University of Oxford and Nordic Mensa Fund.

MLK is supported by Center for Music in the Brain and Centre for Eudaimonia and Human Flourishing, which is funded by the Pettit and Carlsberg Foundations.

## Author contributions

LB, GFR and PV conceived the hypotheses. LB designed the study. LB, MLK and PV recruited the resources for the experiment. LB, GFR and FC collected the data. LB, GFR, MR and AG performed pre-processing, computed BROAD-NESS and the statistical analysis. MLK, GFR, MR, AG and PV provided essential help to interpret and frame the results within the neuroscientific literature. LB, AM and GFR wrote the first draft of the manuscript. GFR and LB prepared the figures. All the authors contributed to and approved the final version of the manuscript.

